# Effect of Human Mesenchymal Stem Cells on Freund’s adjuvant-induced Rheumatoid Arthritis in Sprague Dawley Rats

**DOI:** 10.1101/2021.12.20.473415

**Authors:** Satyen Sanghavi, Vinayak Kedage, Rajesh Pratap Singh, Parvathi Chandran, Vidya Jadhav, Sujata Shinde

## Abstract

**Introduction:** Mesenchymal stem cells (MSC) therapy is a new approach to treat RA. Studies evaluating anti-inflammatory effects of MSCs per RA severity are scarce. Our primary objective was to evaluate anti-inflammatory effects, change in cytokine levels and cartilage regeneration of two different MSC preparations delivered through two different routes of administration in three RA stages: mild, moderate and severe.

**Methods:** Human-derived umbilical cord tissue MSCs (hUCT-MSCs) and human bone marrow-derived MSCs (hBM-MSCs) delivered via intra-plantar and intravenous routes were tested in Freund’s adjuvant-induced arthritis in rats. Arthritis severity was based on the arthritis score (<3=mild, 3=moderate and 4=severe). Assessments included changes in arthritis scoring, paw swelling, haematology parameters, biomarkers (TNFα and IL-10) and histopathology analysis.

**Results:** MSC treatment significantly reduced arthritis scores in all treatment groups. IL-10 levels increased 30 days after treatment with (IP)hUCT-MSCs (*P*=0.0241), (IV)hUCT-MSCs (*P*=0.0095) and (IP)hBM-MSCs (*P*=0.0002). TNF-α levels reduced compared to positive control at 30 days: (IP)hUCT-MSCs (*P*=0.0060), (IV)hUCT-MSCs (*P*=0.0003), (IP)hBM-MSCs (*P*=0.0005), (IV)hBM-MSCs (*P*<0.0001) and continued through 30-60 days. Microscopic examination showed regenerative changes in animal joints treated with both intra-plantar or intravenous MSCs. Arthritis scores reduced in all RA severity groups while benefits (changes in IL-10 and TNF-alpha) were more pronounced in moderate and severe RA. Haematology parameters remained similar among all animal groups at baseline, 30 days and 60 days indicating safety of MSCs.

**Conclusion:** Treatment with hUCT MSCs and hBM MSCs were safe, well-tolerated and effectively reduced joint inflammation, synovial cellularity and pro-inflammatory cytokine levels in CFA-induced RA rat model.

## INTRODUCTION

Rheumatoid arthritis (RA) is a systemic autoimmune disease that causes inflammation of the joints. The global prevalence of RA is estimated to be 460 per 100 000 population between 1980 and 2019.[1] It primarily affects the small joints of the hands and feet. In addition to inflammation in a joint lining called synovium, the aggressive front of tissue (called pannus) invades and destroys local articular structures. RA remains a multifactorial disease involving several genetic and environmental factors.[2] The pathophysiology of RA is poorly recognized, and therefore despite recent therapeutic advances, no definite cure for RA exists.[3-5]

A combined effect of genetic and environmental factors is suggested to trigger development of RA. The contribution of genetic factors is mainly attributed to certain human leukocyte antigen (HLA) alleles of major histocompatibility complex (MHC) class II.[6] Citrullination of peptides post-translationally leads to binding to the shared epitope (SE) with high affinity. This initiates citrulline-specific T- and B-cell responses. Various citrullinated peptides, including fibrinogen, vimentin, a-enolase, aggrecan, and type II collagen (CII) are found in RA joints and are targeted by lymphocytes in RA patients.[6] Additionally, activation of autoreactive CD4+T cells lead to differentiation of CD4+T cells into inflammatory T helper (Th) cell subsets, which produces either interferon (IFN)-γ and tumor necrosis factor (TNF) (Th1), interleukin (IL)-17, IL-21 (Th17) or a mixed cytokine profile (Th1/17) that accumulates in the inflamed joint.[6] These drive differentiation of B lymphocytes into plasma cells, which produce autoantibodies such as anti-citrullinated protein antibodies ACPAs. This promotes osteoclast differentiation and activation, leading to cartilage and bone erosion.[6]

The conventional treatment regimen is a progression through corticosteroids, non-steroidal anti-inflammatory drugs (NSAIDs), non-biologic disease-modifying anti-rheumatic drugs (DMARDs), and biologic DMARDs (e.g., anti-tumor necrosis factor (anti-TNF) and Janus kinase (JAK) pathway inhibitors).[7, 8] These therapies primarily target suppression of inflammatory cascade with limited or no success in controlling the progression of bone and cartilage destruction.[9] Thus, 15–40% of RA patients become resistant to long-term treatments and can become non-responsive to all existing clinical therapies indicating a need for novel RA therapies.[10-13]

Cell therapy with mesenchymal stem/stromal cells (MSCs) is a new approach to treat RA. Due to its dual properties of immunomodulation and tissue repair, mesenchymal stem cells (MSCs) have shown promising results in various autoimmune and degenerative diseases.[14] Several recent studies have shown that MSCs derived from different sources regulate the immune response, including *in vitro* inhibition of T cell proliferation, B cell function, and dendritic cell maturation; however, the underlying mechanisms are not fully elucidated.[15, 16] A couple of systematic reviews of preclinical studies have demonstrated the potential of MSCs in the treatment of RA.[17, 18] Since bone marrow (BM) was the first tissue source used to isolate and expand MSCs (BM-MSCs) for cell-based therapies, BM-MSCs remain the most commonly used cells in animal models of RA.[19] Due to the limitations associated with the BM-MSCs collection and yield, alternative tissue sources of MSCs are also subjected to research, and among this, the umbilical cord is most frequently used. These cells have been administered by a different route, but in general, MSC-based therapy has shown similar beneficial effects independent of the route of administration.[18] A systematic review showed improved efficacy of MSCs therapy when it was administered during the early phases of the disease (47% and 25% in the onset and the established phase of the disease, respectively) rather than at pre-onset (19%) or during chronic phases (9%) of the disease.

Conversely, the current RA clinical trials with MSC-based therapy have been conducted in very refractory RA patients with a long history of the disease, which may have hindered the beneficial effects of MSC-based therapy in the clinic. There is a lack of studies evaluating the anti-inflammatory effects of MSCs per the severity of RA. Various preclinical studies have used the visual scoring system (scores) to evaluate disease severity.[20] With this background, we investigated the anti-inflammatory potential of human-derived umbilical cord tissue MSCs (hUCT-MSCs) and human bone marrow-derived MSCs (hBM-MSCs) in experimental CFA-induced arthritis in Sprague-Dawley (SD) rats. The primary objective of the study was to evaluate the anti-inflammatory effects, changes in the cytokine balance, and cartilage regeneration after administration of these two different MSCs preparation delivered through two different routes of administration in three RA stages: mild, moderate, and severe.

## MATERIALS AND METHODS

### Animals and Ethics

This study was conducted in the facility of Preclinical Research and Development Organization (PRADO), Private Limited, Pune. Due to COVID-19, a lockdown was imposed in India within the study duration. The approval was taken from the IC-SCR committee to initiate this study, including the usage of stem cells in animals (RBPL/IC-SCR/20/002). All procedures followed for the conduct of this study were in accordance with the guidelines set by the Committee for the Purpose of Control and Supervision of Experiments on Animals (CPCSEA). Prior approval of the Institutional Animal Ethics Committee (IAEC) was also obtained IAEC-20-018.

A Sprague Dawley (SD) rat model was selected for this study based on the available literature; it is a widely used model for evaluating the efficacy of the chemicals and for the evaluation of RA.[21-23] A total of 75 Sprague Dawley rats were used in this study. All rats were acclimatized for a period of nine days prior to test item administration. The age of all animals was 6-12 weeks with 202.0–214.5 gm body weight. The atmospheric condition was maintained at temperature 19^°^ to 25^°^C and humidity 30% to 70% throughout the study period. Standard rodent diet and reverse osmosis water treated with ultraviolet light were provided *ad libitum* with a reversed dark-light cycle. During the period of 60 days, all animals were observed once a day for their clinical signs and twice a day for their mortality.

Animals were randomly divided into seven groups; normal control, hUCT-MSCs via i.p. ([IP] hUCT-MSCs) route at hind paws (paw region), hUCT-MSCs via i.v. through tail vein ([IV] hUCT-MSCs) route, hBM-MSCs via i.p. ([IP] hBM-MSCs) route, hBM-MSCs via i.v. ([IV] hBM-MSCs) route, positive control, and methotrexate (MTX) reference group. Adjuvant induced arthritis (AIA) was induced by a single subcutaneous (SC) injection of 0.1 ml of complete Freund’s adjuvant (CFA) (SIGMA-ALDRICH, Lot no. SLCB4664, Product code. 1002865531) containing 10 mg/ml of heat-killed Mycobacterium tuberculosis in all animals except those in the normal control group. All animals were monitored for the development of arthritis for a period of 10 days. Animals from the normal control group received only normal saline solution by intravenous route. Based on the severity score of arthritis, animals were further divided into mild, moderate, and severe subgroups for each group except the normal control group. To induce severe inflammation, the second dose of CFA was administered in moderate and severe groups after 12 days of the first CFA dose.

All animals from (IP)hUCT-MSCs, (IV)hUCT-MSCs, (IP)hBM-MSCs, and (IV)hBM-MSCs groups received the treatment of test items on day 1 and day 10 post-arthritis induction. Six animals each from (IP)hUCT-MSCs, (IV)hUCT-MSCs, (IP)hBM-MSCs, and (IV)hBM-MSCs groups, having arthritis score 2 = mild and three animals each from (IP)hUCT-MSCs, (IV)hUCT-MSCs, (IP)hBM-MSCs, and (IV)hBM-MSCs groups with arthritis score 3 = moderate arthritis and 4 = severe were injected with two test products. Animals from the positive control group did not receive any treatment and were considered as the positive control. Animals in the methotrexate (MTX) reference group and having arthritis score 2 or 3 received Methotrexate (MTX) orally (p.o.) at 7 mg/kg once a week for 5 weeks in mild and 9 weeks in moderate RA group. Analysis of some efficacy parameters in the Methotrexate group was unavailable due to manpower issues to handle/look after animals during the imposed lockdown due to COVID-19 pandemic.

### Test drug information

Regrow Biosciences Pvt. Ltd., Nangargaon, Lonavala, has manufactured hUCT-MSCs and hBM-MSCs for this study. All animals of treatment groups ((IP)hUCT-MSCs, (IV)hUCT-MSCs, (IP)hBM-MSCs, and (IV)hBM-MSCs) were administered a cell suspension containing 4×10^6^ cells and 1.5×10^6^ cells via the intravenous and intra-plantar route, respectively. As per the literature available for the selection of the desired dose to be injected into the SD rats, MSCs doses starting from 1 million to 5 million were used in different studies.[24-28]

### Efficacy parameters

#### Arthritis Scoring or Paw Swelling

Development of arthritis in hind paw of animals was scored on a scale of 0–4, where 0 = no swelling, 1 = minimal or slight swelling and erythema of the limb, 2 = mild swelling and erythema of the limb, 3 = moderate or gross swelling and erythema of the limb, and 4 = severe swelling (figure 1). The scoring was performed by visual observations by two different technicians to minimize the error in the paw swelling score.

**Figure 1.**
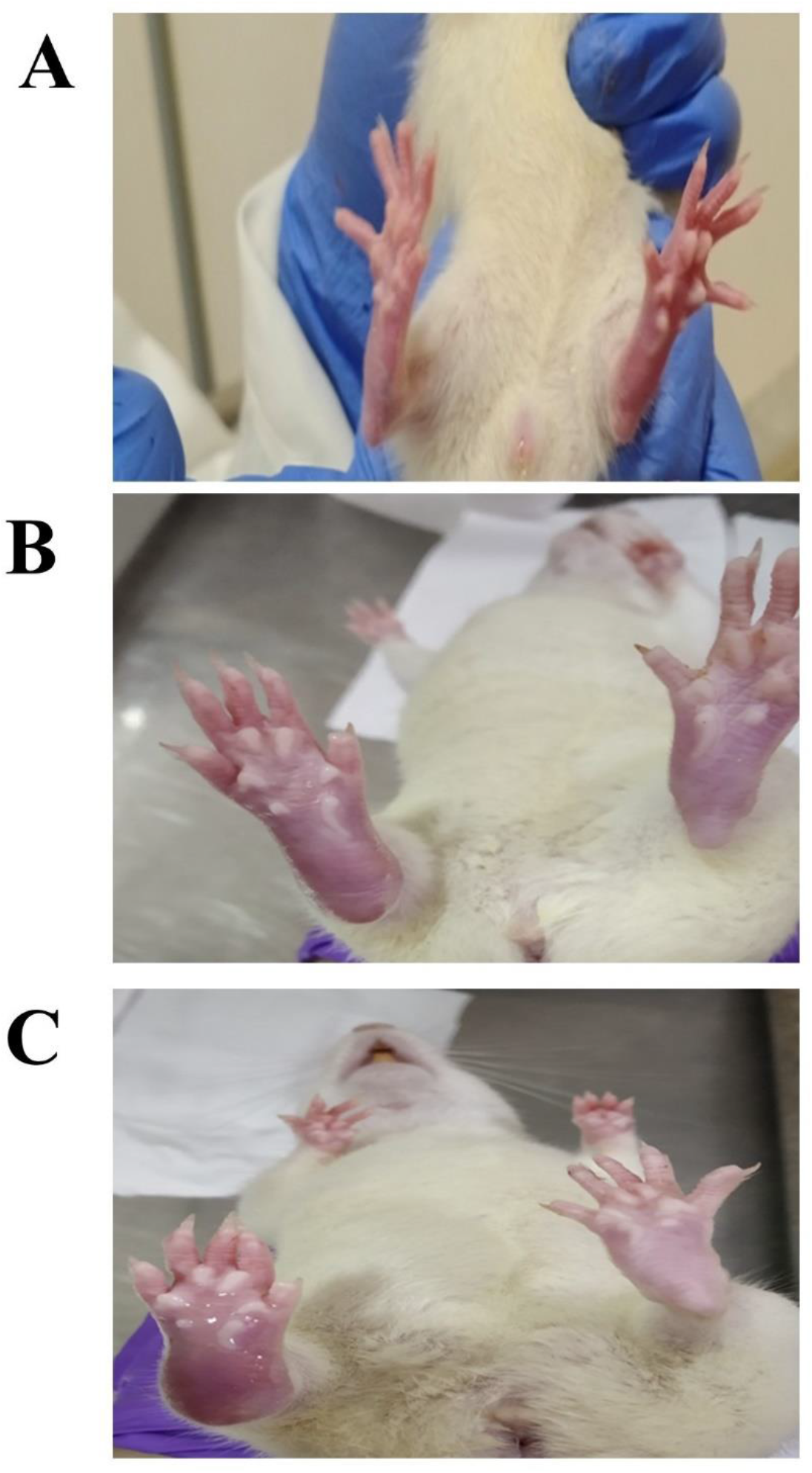
Development of rheumatoid arthritis (RA) model with different severity (A) mild RA (B) moderate RA (C) severe RA

#### Hematology

After completion of the observation period, animal blood samples from all treatment and control groups were collected from retro-orbital sinuses for evaluation of hemoglobin content (HGB), total red blood cell count (RBCs), % packed cell volume (% PCV), total white blood cell count (WBCs) and erythrocyte sedimentation rate (ESR).

#### Histopathology

After blood collection, animals were euthanized by CO_2_ asphyxiation; hind paws were collected in 10% neutral buffered formalin, decalcified, and subjected to routine histopathological examination stained with Hematoxylin and Eosin stain.

#### Biomarkers Analysis

Serum collected from each animal were separated, and samples were stored at −20^°^C and used for cytokines estimation viz., TNFα (Make: Thermofischer; Catalogue no.: KRC3011), IL-10 (Make: Thermofischer; Catalogue no.: BMS629) by ELISA method as per the standard protocol provided by the manufacturer.

### Safety parameters

After dosing the test item, all the animals were observed carefully once daily for treatment-related clinical signs and twice daily for morbidity/mortality. The body weight of all the animals was recorded on a weekly basis.

### Statistical analysis

Results are expressed as mean±standard deviation (SD). Statistical significance of difference was determined using the ANOVA test for comparison between different groups. The SAS^®^ version 9.3 was used for all statistical analyses. *P*<0.05 was considered statistically significant.

## RESULTS

### Effect on Rheumatoid arthritis score

As shown in figure 2, there was a significant reduction in the arthritis scores after 30 days of treatment with MSCs in all four groups (*P*<0.0001 for all four group’s comparison with positive control) and also after the treatment with the methotrexate (*P*<0.0001 vs. positive control). Treatment with MSCs showed a significant reduction in arthritis scores in all three RA severity groups, mild, moderate, and severe.

**Figure 2.**
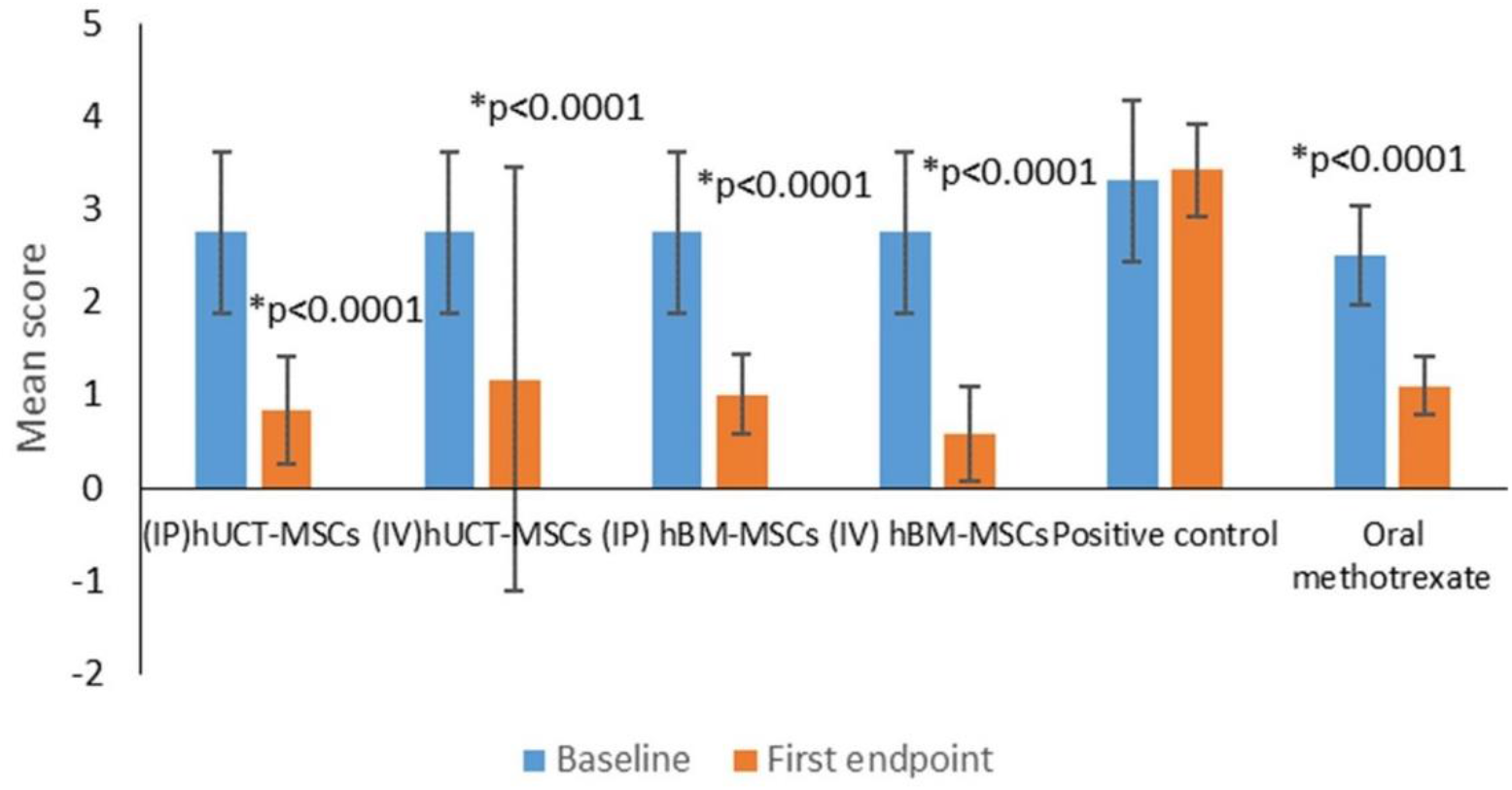
Changes in rheumatoid arthritis (RA) scores in animals after treatment. Only statistically significant p values (p<0.05) are presented in the graph * vs positive control at first endpoint. IP, Intraperitoneal; IV, intravenous; hUCT-MSCs, human-derived umbilical cord tissue MSCs; hBM-MSCs, human bone marrow-derived MSCs.

### Effect on IL-10 levels

IL-10 is an anti-inflammatory interleukin leukotriene. After ten days of CFA administration, there was a statistically significant reduction in the IL-10 levels in all the groups (*P*<0.0001). Treatment with MSCs did not show effect at 30 days, but the IL-10 levels started increasing after 30 days that reached a statistically significant increase in animals in (IP)hUCT-MSCs (*P*=0.0241), (IV)hUCT-MSCs (*P*=0.0095), (IP)hBM-MSCs (*P*=0.0002) groups compared with IL-10 levels in the positive control (figure 3A). When the changes in IL-10 levels were analyzed by the severity of the disease, effects on IL-10 levels were more pronounced in animals with moderate and severe RA (figures 3B and 3C).

**Figure 3.**
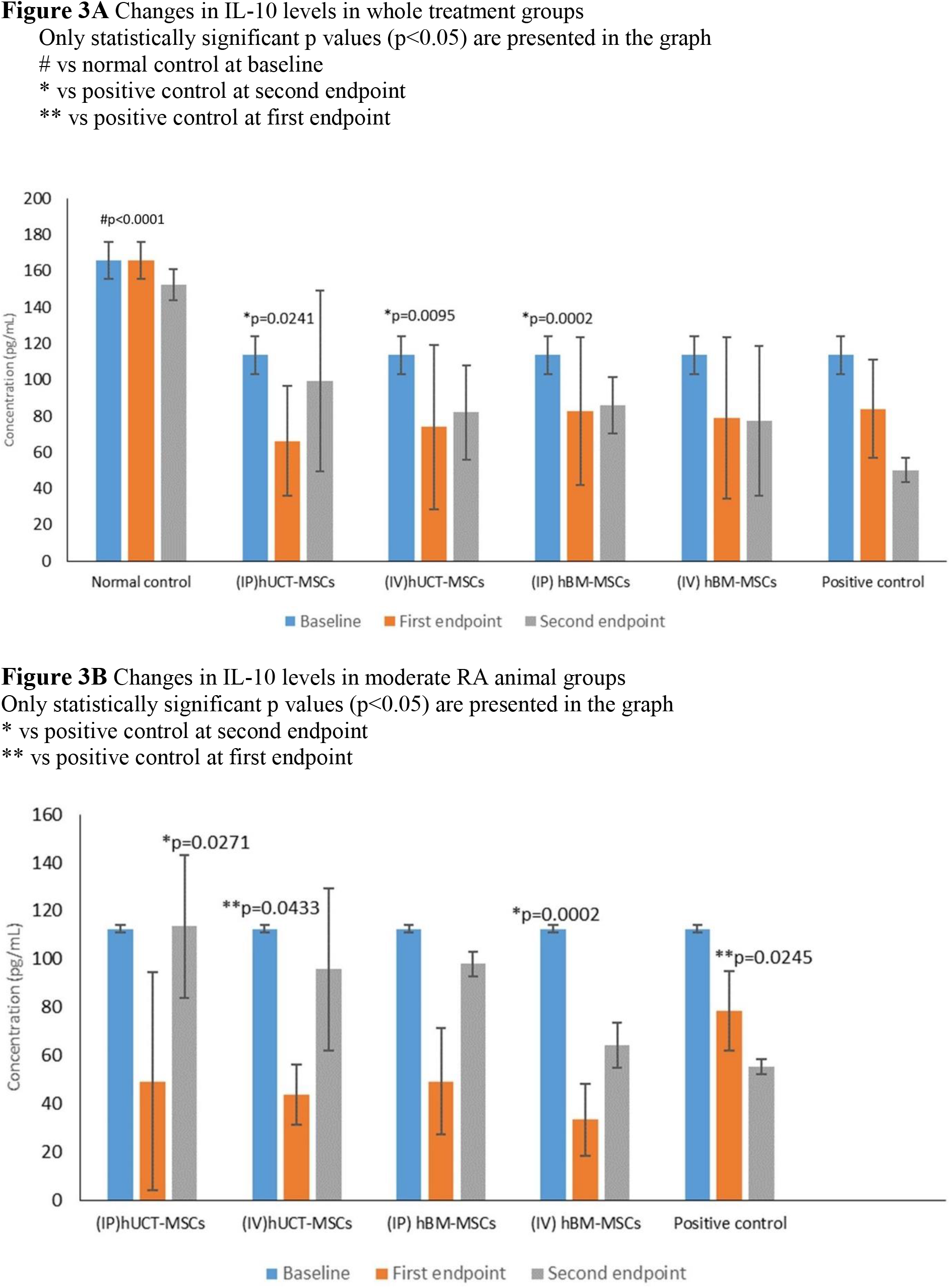
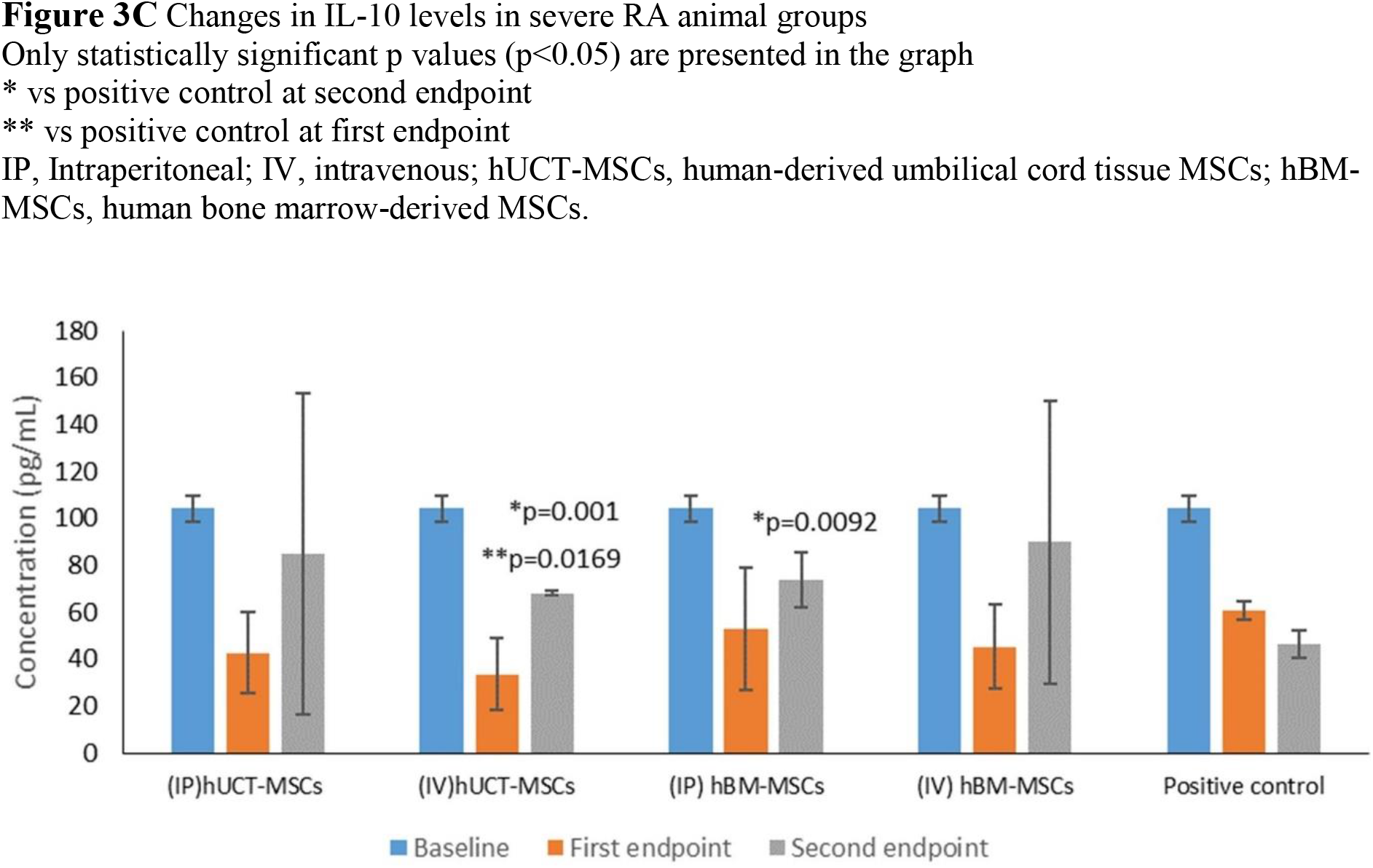
Changes in interleukin 10 (IL-10) levels in animals after treatment.

### Effects on TNF-alpha levels

TNF-alpha is an inflammatory mediator, levels of which were increased significantly following the CFA administration in positive control animals compared with the normal control (*P*<0.0001) at baseline (figure 4A to 4C). As shown in figure 4A, treatment with MSCs showed statistically significant reversal of this increase compared with positive control at 30 days: (IP)hUCT-MSCs (*P*=0.0060), (IV)hUCT-MSCs (*P*=0.0003), (IP)hBM-MSCs (*P*=0.0005), (IV)hBM-MSCs (*P*<0.0001). The trend of decrease in TNF-alpha levels continued even through 30-60 days when compared with positive control with statistically significant difference: (IP)hUCT-MSCs (*P*=0.0029), (IV)hUCT-MSCs (*P*=0.0037), (IP)hBM-MSCs (*P*=0.0043), (IV)hBM-MSCs (*P*=0.0048).

**Figure 4.**
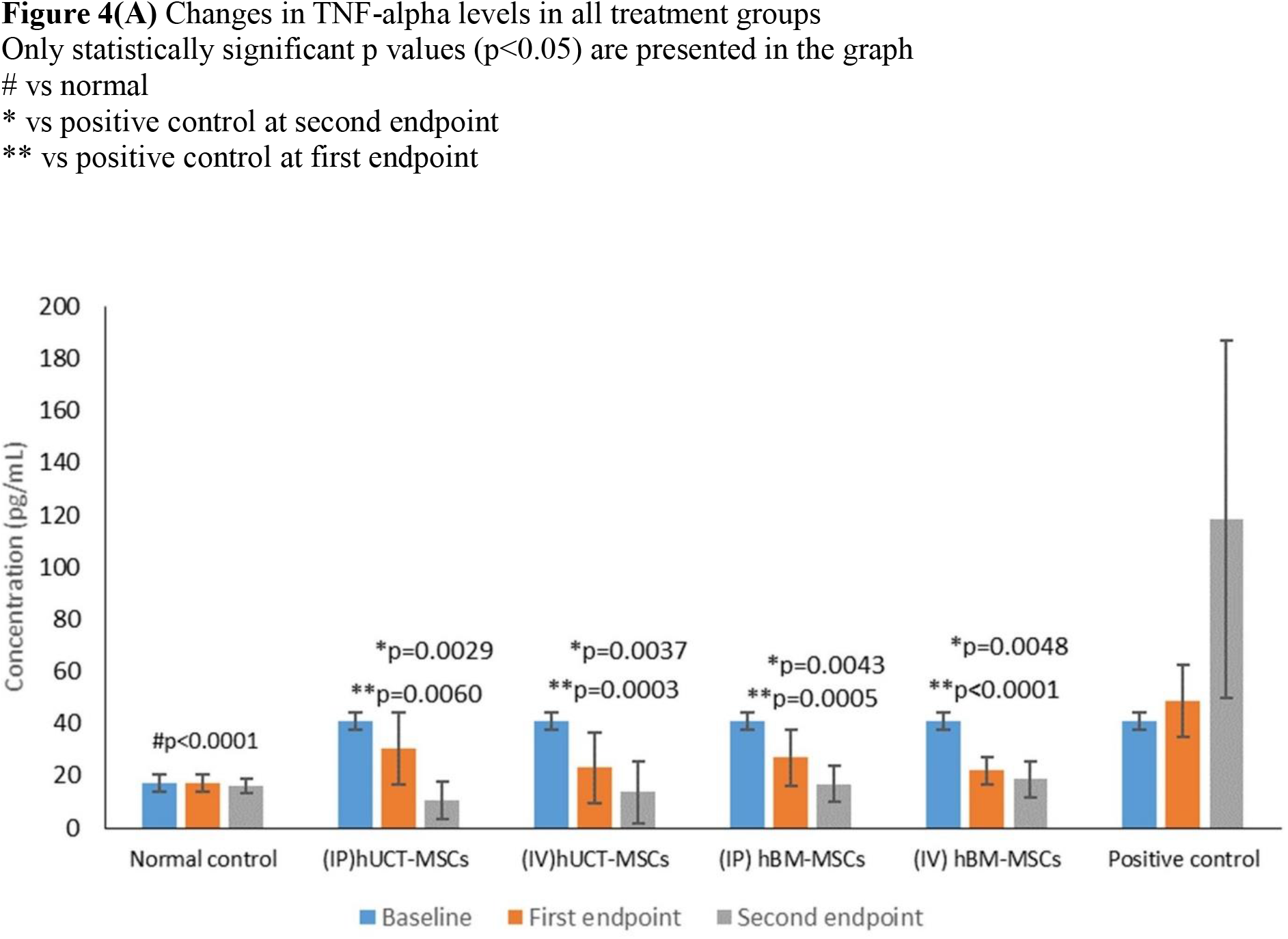
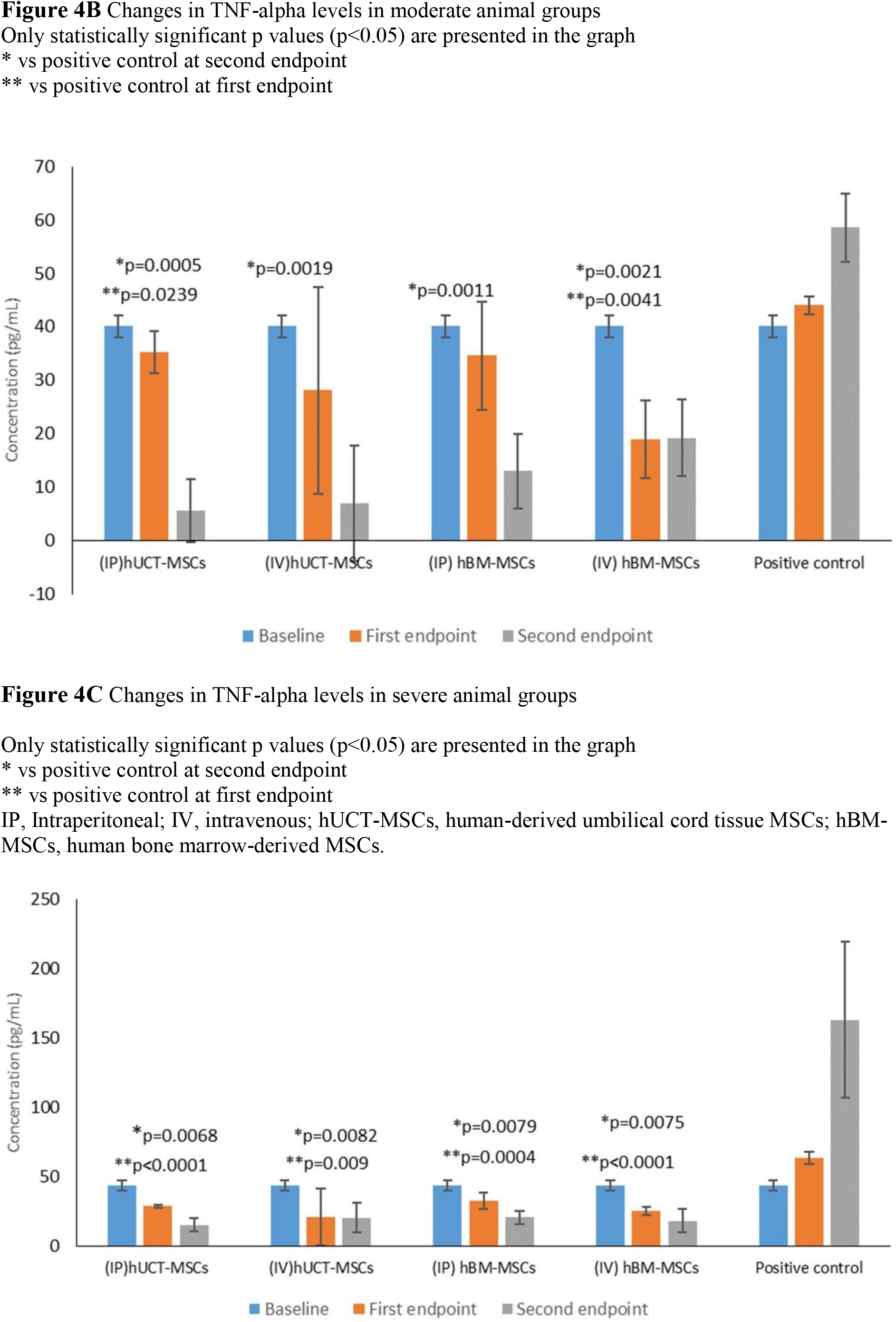
Changes in tumor necrosis factor alpha (TNF-alpha) levels in animals after treatment.

When the changes in TNF-alpha levels were analyzed by the severity of the disease, effects on TNF-alpha levels were more pronounced in animals with severe RA (figures 4C), with all the treatment groups showing a significant change in TNF-alpha levels starting from 30 days and continuing until 60 days. Whereas in the animals with moderate RA, only MSCs treatment groups had shown the change in TNF-alpha levels at 30 days (figure 4B).

### Changes in the histopathological characteristics

Microscopic examination of animals’ joints from hUCT-MSCs groups (figure 5) and hBM-MSCs groups (figure 5) showed regenerative changes when administered either by intra-plantar or by intravenous routes. However, the incidence of changes such as uniformity in the chondrocyte layer and presence of zones of articular cartilage was better in animals receiving test items by intravenous route than in animals receiving them via the intra-plantar route.

**Figure 5.**
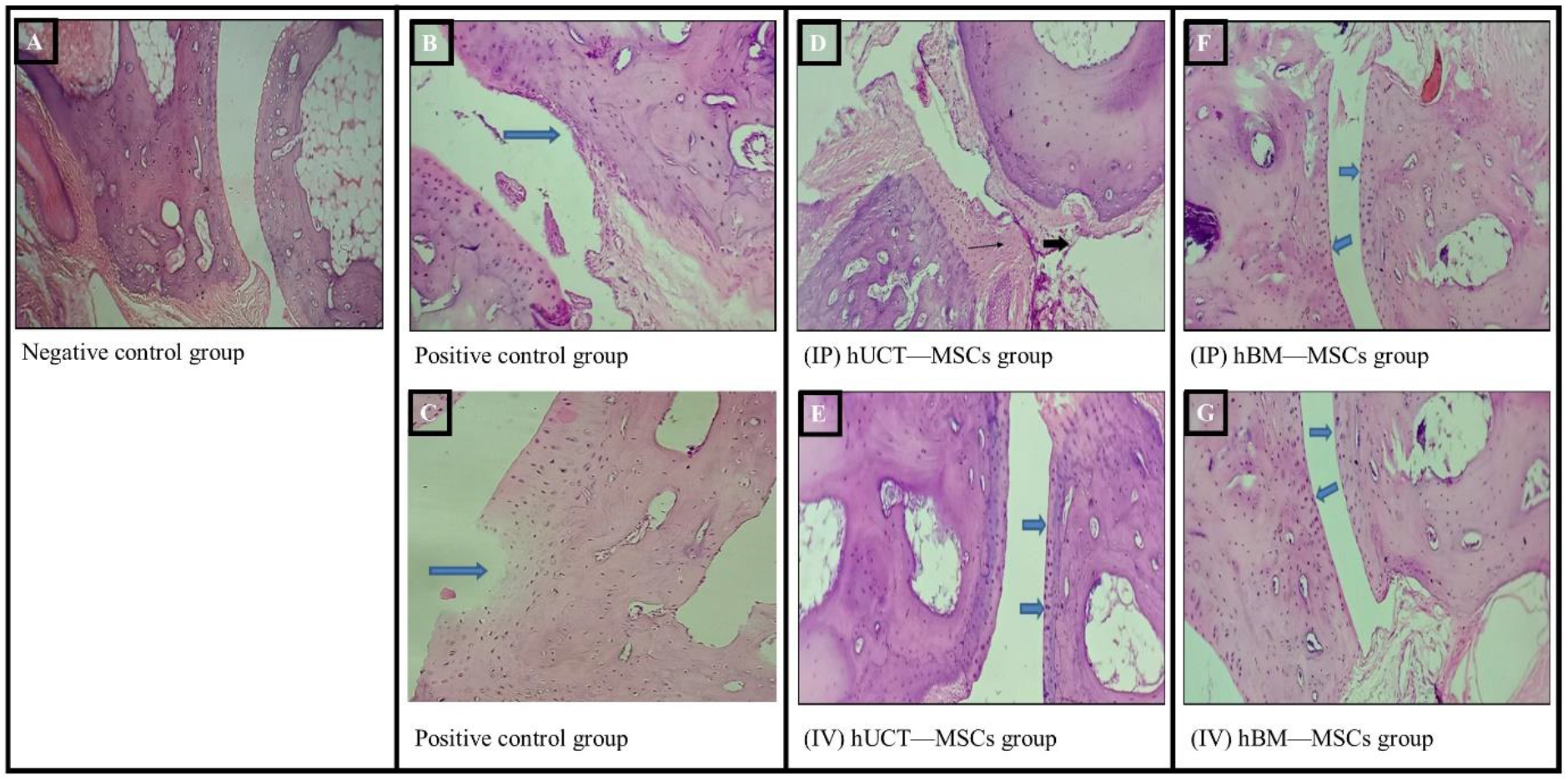
Histological changes in animals after treatment with human umbilical cord tissue mesenchymal stem cells (hUCT-MSCs) and human bone marrow mesenchymal stem cells (hBM-MSCs) via intraplantar (IP) and intravenous (IV) routes, respectively. (A) Normal control group animals showed intact architecture and uniformity of chondrocyte layer, presence of zones of articular cartilage and cell organization (B and C) Positive Control group animals showed severe destruction of bone and cartilage as well as hypercellularity of chondrocyte (indicated by arrow head). H&E 10X (D) (IP) hUCT-MSCs with 1.5 × 10^6^ cells dose showed mild cellular infiltration (arrow) and erosion (arrow head) in synovial tissue of severe group animals (H&E 10X). (E) (IV) hUCT-MSCs with 4 × 10^6^ cells showed minimal destruction in cartilaginous and bone layer (arrow head) with mild mitotic figures (arrow) in severe group animals (H&E 10X). (F) (IP) hBM-MSCs with 1.5 × 10^6^ cells dose showed intact articular cartilage with normal joint cavity and synovial tissue with minimal mitotic figures (arrow head) in severe group animals. H&E 10X. (G) (IV) hBM-MSCs with 4 × 10^6^ cells dose showed intact articular cartilage with normal joint cavity and synovial tissue in severe group animals (H&E 10X).

### Changes in hematology parameters

As seen in figure 6, the WBCs, RBCs, HBG, and ESR concentrations were similar among the animal groups at baseline, 30 days, and 60 days (*P*>0.05). This indicated that treatment with MSCs was safe and well-tolerated. No infection, abnormality, and mortality, were observed.

**Figure 6.**
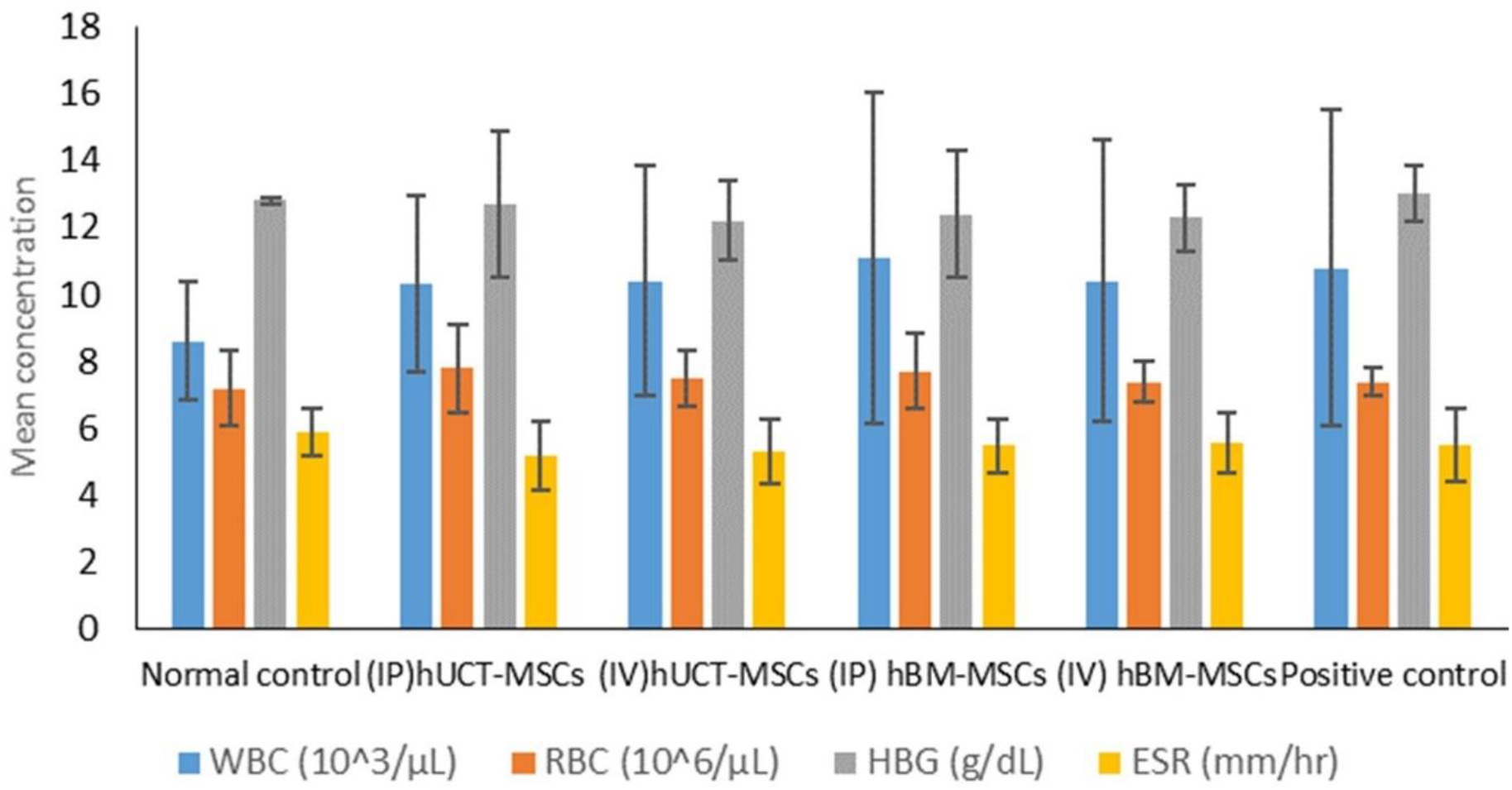
Changes in hematology parameters in animals after treatment. WBC, white blood cell count; RBC, red blood cell count; HBG, hemoglobin content; ESR, erythrocyte sedimentation rate; IP, Intraperitoneal; IV, intravenous; hUCT-MSCs, human-derived umbilical cord tissue MSCs; hBM-MSCs, human bone marrow-derived MSCs.

## DISCUSSION

MSCs have shown promising potential in treating several autoimmune diseases, including RA.[15-17] In the present study, we demonstrated the effect of MSCs at various stages of RA using the CFA-induced RA rat model. Our results showed that the signs and symptoms of RA improved 30 days following the treatment with MSCs, as shown by the decrease in arthritis score. The favorable changes in IL-10 and TNF-alpha were noted after treatment with MSCs, which were more pronounced in moderate and severe RA compared to animals in mild groups. Histopathological changes showed regeneration of synovial membrane cells and articular cartilage. Our findings reiterated the therapeutic potential of MSCs for the treatment of RA.

RA in humans is a systemic chronic autoimmune disease affecting multiple organs in the body, causing predominantly inflammatory synovitis and structural damage to bone and cartilage.[29] Although animal models cannot replicate the exact clinical manifestations of RA in humans, several animal models have been used to understand possible treatments for RA.[30] Several studies indicated that MSCs possess an immunosuppressive role, with potential in the treatment of RA.[31-36] However, there remain uncertainties about the efficacy of this therapy regarding different MSC sources (bone marrow, adipose tissue, umbilical cord, MSC cell lines), doses (0.5–5×10^6^, single or multiple infusions), routes of administration (intravenous, intraperitoneal, intra-articular), the timing of delivery (before disease onset, early or advanced disease), MHC match (syngeneic, allogeneic, xenogeneic) used in literature studies.

In this study, we sought to investigate the anti-arthritis effects of MSCs from two different sources (human umbilical cord blood and human bone marrow) administered through two different routes of administration (intra-plantar and intravenous). Both treatments increased IL-10 concentration and reduced the TNF-alpha levels even in moderate to severe disease groups. This was also correlated with the observed decrease in arthritis score in all the MSCs treatment groups compared with the control group. Histopathological examination with H and E stain revealed that tissue damage caused by CFA administration was repaired via the newly regenerated cartilage by treatment with both types of MSCs, even in animals with moderate to severe disease in this study. The trends of change in the IL-10 and TNF-alpha levels were similar to previous studies.[25, 37, 38] Favorable histopathological characteristics like regeneration of synovial membrane cells, normal articular cartilage, and normal synovial membrane have been seen in previous studies.[23, 32, 39]

A recent meta-analysis of preclinical studies showed that MSC-based therapy modulates and inhibits arthritis inflammation regardless of the tissue source and route of administration in the majority of preclinical animal studies.[19] Results from our study were in line with this meta-analysis, providing a rationale for using MSC-based therapy in the clinic. As a result of the encouraging preclinical data, there has been a rapid rise in the use of MSC-based therapy in clinical trials of RA recently; however, these studies have mainly involved very refractory RA patients. Data from the preclinical studies suggest that the early phases of inflammation are best suitable for MSC infusion in contrast to most of the completed clinical trials with MSC based therapy that have been conducted in RA patients with a long history of the disease, which could have masked the clinical benefit of the MSC-based therapy. Interestingly, RA patients during the early phases of the disease are now being recruited in ongoing clinical trials. This change in the inclusion criteria may increase the efficacy of MSC-based therapy.[40] Furthermore, a better understanding of the pathogenesis of the disease and the mechanisms of action underlying MSC therapy in RA would identify key biomarkers and contribute to the deployment of optimized protocols of MSC-based therapy for the benefit of RA patients with unmet medical needs.

Despite using DMARDs and novel biological agents, there is still no promising cure for RA. Almost 50% of patients with RA have experienced remission, and 10%–15% have been found refractory or develop severe adverse effects to the treatments and cartilage degeneration.[18] Conversely, in our study, we could observe cartilage regeneration using MSCs in the RA animal model. We also observed a decrease in TNF levels showing better outcomes with MSCs treatment. The development in this area may lead to a cellular therapy that will be patient-friendly, can be given at home as IV infusion 1-3 times a year, and will be a paradigm shift in the management of RA patients. However, well-designed studies are required to validate its effect in further preclinical studies and its impact on patient life, activity, career, steroid-led complications, treatment cost, and frequent doctor visits in future clinical studies.

Our results show that treatment with both types of MSCs viz., hUCT MSCs and hBM MSCs effectively reduced joint inflammation, synovial cellularity, and the levels of pro-inflammatory cytokines in the CFA-induced rheumatoid arthritis model in rats. The treatment with test products ameliorated histopathological changes in joints caused by administration of CFA and modulated the expression of cytokines. The treatment was safe, as indicated by no change in the hematological parameters. However, the results need to be substantiated in well-designed clinical studies.

## Acknowledgment

Authors would like to thank Prado Preclinical Research Pvt. Ltd. for providing support in conducting this animal study. Authors also thank CBCC Global Research for providing medical writing support in development of this manuscript.

## Contributors

SS, VK, VJ, RS, SS and PC contributed to the conception and study design. VK, VJ, RS, SS and PC acquired and analysed the data. SS, VK, VJ, RS, SS and PC contributed to interpretation of the data. All authors wrote the first version of the manuscript and revised it critically. All authors read and approved the final manuscript.

## Funding

This study was funded by Regrow Biosciences Pvt. Ltd.

## Conflict of interest

Satyen Sanghavi, Dr. Vinayak Kedage, Rajesh Pratap Singh, Parvathi Chandran, Vidhya Jadhav and Sujata Shinde are employees of Regrow Biosciences Pvt. Ltd.

## Patient consent for publication

Not required.

## Ethics approval

The study was approved by the Institutional Animal Ethics Committee (IAEC) IAEC-20-018.

## Data sharing statement

All data relevant to the study are included in the article or uploaded as supplementary information.

